# Pollination responses to artificial light at night may vary with lighting arrangement

**DOI:** 10.64898/2026.07.10.737511

**Authors:** Dongbo Li, Lily A.J. Adeniji, Rochelle J. Meah, Christopher F. Clements

## Abstract

Artificial light at night (ALAN) can alter the movement of nocturnal pollinators such as moths, with potential consequences for pollination services. However, most studies have focused on the presence or properties of lights, while the role of spatial lighting configuration remains poorly understood. We conducted a small-scale field experiment to test how different lighting configurations affect pollination success in moth-pollinated plants, using low-intensity LED garden lights. Potted phytometer plants of three moth-pollinated species were exposed to one of three treatments: no-light control, isolated LED point lights, or 25 m linear arrays of multiple LED lights. We quantified pollination success as both the probability of seed set and reproductive output through seed and capsule production. Experimental lighting affected plant reproductive success, with light arrays increasing seed-set probability, seed and seed capsule number relative to unlit controls, whereas point lights showed no clear effect. These results provide preliminary evidence that the spatial configuration of artificial lights may influence nocturnal pollination outcomes. Future work combining phytometer assays with direct tracking of moth movement is needed to assess whether light arrays facilitate or redirect the dispersal of nocturnal pollinators.

## 1. Introduction

Artificial light at night (ALAN) is rapidly expanding worldwide and increasingly recognized as an emerging driver of environmental change (Gaston & Sánchez de Miguel, 2022; Sanders et al., 2021). By altering the behaviour and distribution of nocturnal pollinators, particularly moths, ALAN can disrupt plant-pollinator networks with cascading consequences for pollination services (Giavi et al., 2021; Knop et al., 2017). Although the direct ecological impacts of ALAN have been well documented (Giavi et al., 2021; Macgregor et al., 2017; Shivanna, 2022), little is known about how the spatial configuration of artificial lights affects the availability of nocturnal pollinators and shapes plant-pollinator interactions. At the landscape scale, lighting configuration may structure pollinator movement paths (Morrell et al., 2024), creating spatial heterogeneity in pollen transfer and plant reproductive success. Understanding ALAN in a landscape context is therefore essential for developing conservation strategies that maintain pollination services in fragmented landscapes (Grenis et al., 2023).

Nocturnal insects often respond to ALAN by flying towards illuminated sources or by altering normal flight movements (Meah et al., 2026; Mungee et al., 2025). Such responses vary with light properties, including light type (Macgregor et al., 2019), spectral composition (Boom et al., 2020; Fabusova et al., 2024), and distance from the light source (Giavi et al., 2020). However, the ecological effects of ALAN may also depend on how light sources are arranged in space. Artificial lights can act as spatial nodes of attraction or repellence within otherwise dark landscapes, and their effects may differ depending on whether they occur as isolated point sources or as arrays. Isolated point lights, such as street lights at road corners, may concentrate night-active insects locally and limit their dispersal across landscapes (Degen et al., 2016). In contrast, arrays of lights such as garden or streetlight networks may influence insect movement across broader spatial scales. Recent studies suggest that linear streetlight arrays can alter moth flight behaviour beyond the directly illuminated area and create barrier effects under some conditions (Degen et al., 2024; Giavi et al., 2020). Low-intensity garden lights provide a useful system for testing these configuration effects, as multiple weak light sources can be arranged into linear networks without creating high illumination levels like in streetlights. Whether such configuration-dependent effects on movement translate into changes in pollination services remains poorly understood.

Here, we present a small-scale exploratory field experiment testing whether the spatial arrangement of low-intensity artificial lights can influence pollination success in moth-pollinated plants. Our aim was not to provide a definitive mechanistic test of moth movement, but to assess whether contrasting lighting arrangements produce detectable differences in plant reproductive outcomes under field conditions. We established three lighting treatments in an open meadow: no-light controls, isolated point lights, and 25-m light arrays (Fig. 1). In the light-array treatment, multiple LED lights were spaced at 1-m intervals along a 25-m transect. In the point-light treatment, two single LED light sources were placed 25 m apart, without intervening lights. The no-light control used the same 25-m spatial layout but without artificial illumination. Within each treatment, we placed potted phytometer plants consisting of three moth-pollinated species *Matthiola longipetala, Silene nutans*, and *Zaluzianskya capensis*, and quantified pollination success using their seed set. Potted phytometer provide a useful way to measure pollination outcomes when the direct observation of nocturnal pollinator movement is difficult (Hackett et al., 2024; Li et al., 2024). Plants were placed at either the both ends of the light arrays (light array treatment), or beside the point lights (point light treatment), or the no-light controls. Only one plant of each species was placed at each position, allowing for the measure of direct pollination outcomes potentially from pollinator movement (similar to Li et al., 2026). By placing phytometers at defined positions within each lighting treatment, we assessed whether different lighting configurations altered plant pollination success.

**Figure 1.**
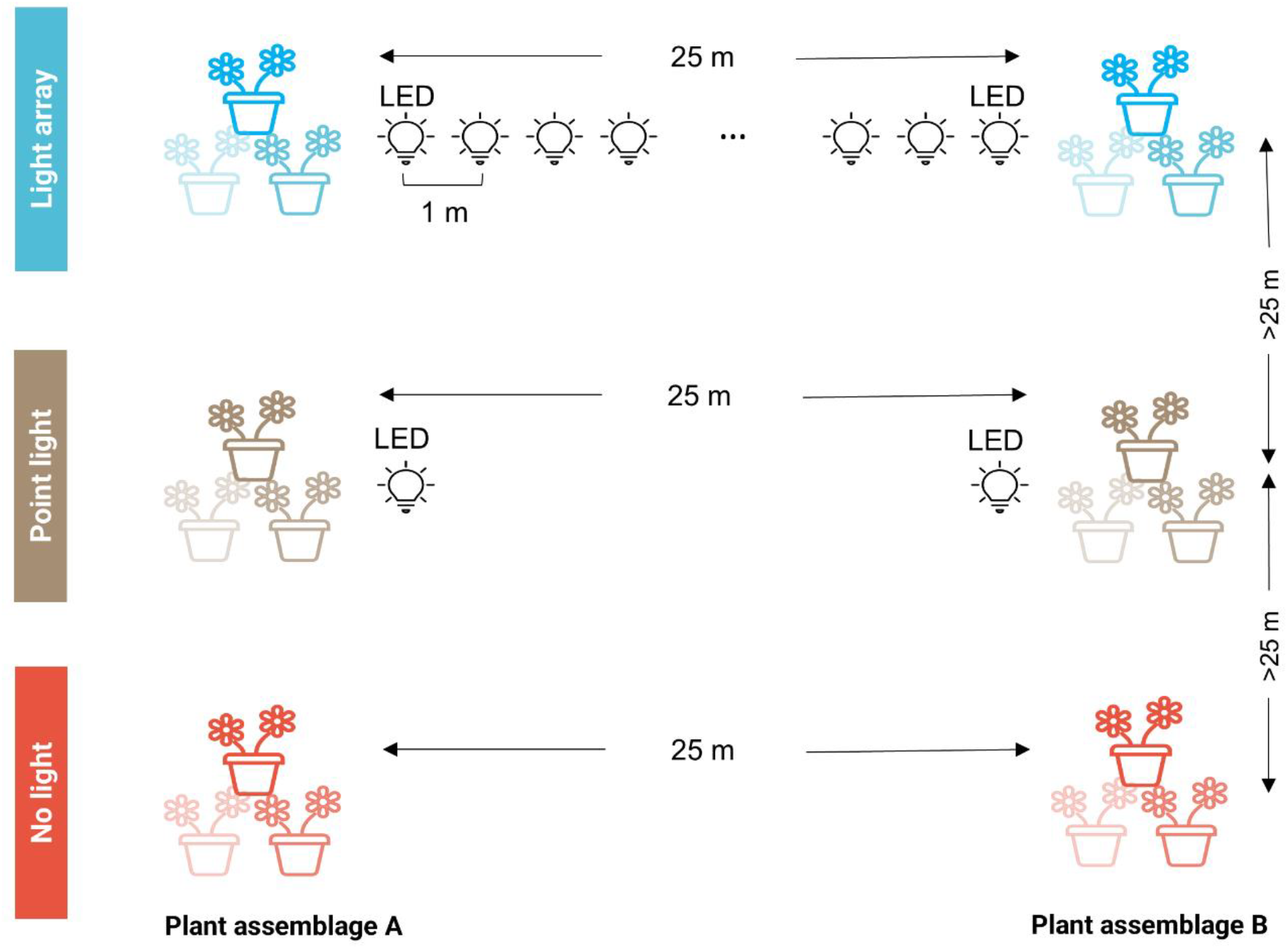
Schematic diagram of the experimental setup. The experiment included three lighting treatments: light arrays, point lights and no light (unlit) controls. Phytometer plant assemblages consisted of one individual of each of three moth-pollinated plant species: *M. longipetala, S. nutans* and *Z. capensis*. Assemblages were placed at either end of each light array, beside each point light, or 25 m apart in no light controls.

## 2. Methods and materials

### 2.1 Study site

The experiment was conducted from June to September 2022 at Fenswood farm, Bristol, United Kingdom (DMS, N 51° 25’ 27”, W 2° 40’ 18”). This site is situated in a rural setting, contains no sources of artificial light and has a minimum distance of 600 m from the nearest village. The site has a relatively low zenith radiance of 0.53 nWcm^-2^sr^-1^ (2025 VIIRS data from NASA’s Black Marble nighttime lights product taken from Jurji Stare, www.lightpollutionmap.info). The farm contains 62 ha of arable farmland, in which nearly two-thirds is intensively farmed with economic crops such as oat and barley, and the rest is used as hayfields and grazing pastures. We selected three hayfields (2.7–5.1 ha) for the placement of the experimental light treatments. All hayfields were surrounded by fens and hedgerows and were largely un-disturbed by anthropogenic activities.

### 2.2 Experimental treatment

Three lighting treatments were established in each hayfield: no-light (unlit) control, point light and light array, giving three replicate blocks per treatment. Each treatment was arranged along a 25 m linear experimental unit, with one endpoint at each end where phytometer plants (see below) were later placed (Fig. 1). In the no-light control, no lights were placed at either endpoint or along the 25 m distance between them. In the point-light treatment, two solar-charged LED lights were placed at each endpoint (two LED lights at one endpoint to reduce the risk of single light failure), with no intervening lights. In the light-array treatment, the two endpoints were connected by a linear array of 26 LED lights: one light at each endpoint and 24 additional lights spaced at 1 m intervals between them. The treatments were designed to represent contrasting field-realistic lighting configurations rather than to standardize cumulative light output across treatments. Point-light treatments represented isolated low-intensity light sources, whereas light-array treatments represented continuous low-intensity lighting networks. Consequently, the array treatment differed from the point-light treatment in both the number of light sources and the spatial extent of illumination, as these are inherent features of linear lighting networks. Lights were solar-charged white LED bollard garden lights (1.2 W, ∼5 lumen, maximum ∼7 lux at 10 cm above ground, GIGALUMI Ltd, China, Fig. S1), positioned approximately 20 cm above ground level and automatically illuminated after dusk for up to 8 h per night (Fig. S2).

The three treatments were arranged in parallel within each hayfield and separated by at least 25 m perpendicular distance. Replicate hayfields were separated by more than 100 m and treated as independent blocks. Lighting treatments remained in place throughout the three-month field season. Treatment positions within each hayfield were randomly reshuffled midway through the field season to reduce potential location effects. Across the three replicate hayfields, the experiment included nine treatment units and 18 endpoint positions for phytomter plants.

### 2.3 Phytometer plants

Three moth-pollinated species, *Matthiola longipetala, Silene nutans*, and *Zaluzianskya capensis*, were used to measure pollination success. Species were selected because of their associations with nocturnal pollinators and their partial or complete dependence on insect pollination. No naturally occurring individuals of any of these species were observed at the study site. *M. longipetala* is a self-incompatible, night-scented species that is particularly attractive to hawkmoths (Kunin & Shmida 1997). Both *S. nutans* and *Z. canpensis* are self-compatible but nocturnal pollination has shown to increase seed set (Johnson et al., 2002; Vanderplanck et al., 2020).

Plants were grown in greenhouse following a standard sowing procedure under pollinator-free conditions. Seeds of each species were sown in 55 × 30 cm plastic trays under uniform lighting conditions (16 h light: 8 h dark) and standard greenhouse temperature (∼18°C). After germination, seedlings were transplanted individually into 7 cm square pots filled with standard multi-purpose compost (Westland Ltd, UK). Potted plants were maintained in the greenhouse until flowering. Once flowering, plants were transferred to the field while remaining in their individual pots. Pots were placed in shallow garden trays to reduce drying during field exposure.

Because the three species differed in flowering phenology under greenhouse conditions, they were deployed in species-specific two-week sampling rounds (Fig. S3). In each round, one flowering potted plant of the focal species was placed at each endpoint of every 25 m experimental unit: at either end of each light array, beside each point-light treatment, or at equivalent positions in the no-light control. Initial plant size was randomized among treatments, and there were no treatment differences in the starting number of flower buds (all *p* > 0.05, Table S1).

Each round of deployment lasted two weeks. *M. longipetala* was deployed over three two-week rounds, giving 54 plants in total, whereas *S. nutans* and *Z. capensis* were each deployed for one two-week round, giving 18 plants per species. The number of plant replication was limited by the logistical demands of moving, watering and keeping potted plants during the field experiment. After field exposure, plants were returned to the greenhouse to allow fruits and seeds to mature. New flower buds produced after field exposure were removed, such that reproductive output reflected flowers exposed during the experimental period. Flower heads were covered with muslin bags to collect seeds while allowing ventilation.

We used the seed set of each plant as a proxy measure of pollination. Seed sets were harvested from plants when they were fully matured. Both seed and capsule numbers of each plant were counted, providing a measure of pollination effect on these phytometer plants.

### 2.4 Statistical analysis

To test whether experimental lighting influenced the probability of seed set, we fitted generalized linear models (GLMs) with a binomial error distribution and logit link. Seed set was coded as a binary response (1 = at least one seed produced; 0 = no seeds produced) per plant. Initial models included treatment (control, point-light, light-array), species identity, and their interaction as a fixed effect. To further test if treatment affects the magnitude of seed set, we then analysed the numbers of seeds and seed capsules produced using negative binomial models with the same explanatory structure. For all models, we compared full interaction and additive models and retained the most parsimonious model based on Akaike’s information criterion (AIC, Table S2). Overall effects were assessed using Type III chi-square tests to each model. Parameter estimates and 95% confidence intervals (CI) of the best fitted models were obtained by non-parametric case-resampling bootstrap procedures (10,000 iterations). Binomial and negative-binomial GLMs were fitted using ‘glmmTMB’ package (Brooks et al., 2017), and post-hoc comparisons and model diagnostic procedures were conducted using the ‘emmeans’ (Lenth, 2025) and ‘DHARMa’ (Hartig & Hartig, 2017) packages in R version 4.5.2 (RCoreTeam, 2025).

## 3. Results

Experimental lighting significantly affected the probability of plants setting seeds (χ^2^ = 10.764, df = 2, *p* = 0.0046, Fig. 2a), with no overall difference among species (χ^2^ = 2.930, df = 2, *p* > 0.05). Relative to unlit controls, light arrays increased the probability of seed set (log-odds = 2.02, 95% confidence interval (CI) = [0.76, 3.74]), whereas point-light treatments did not differ from controls (log-odds = 0.74, 95% CI = [−0.54, 2.20]), nor between point-light and light arrays (*p* > 0.05).

**Figure 2.**
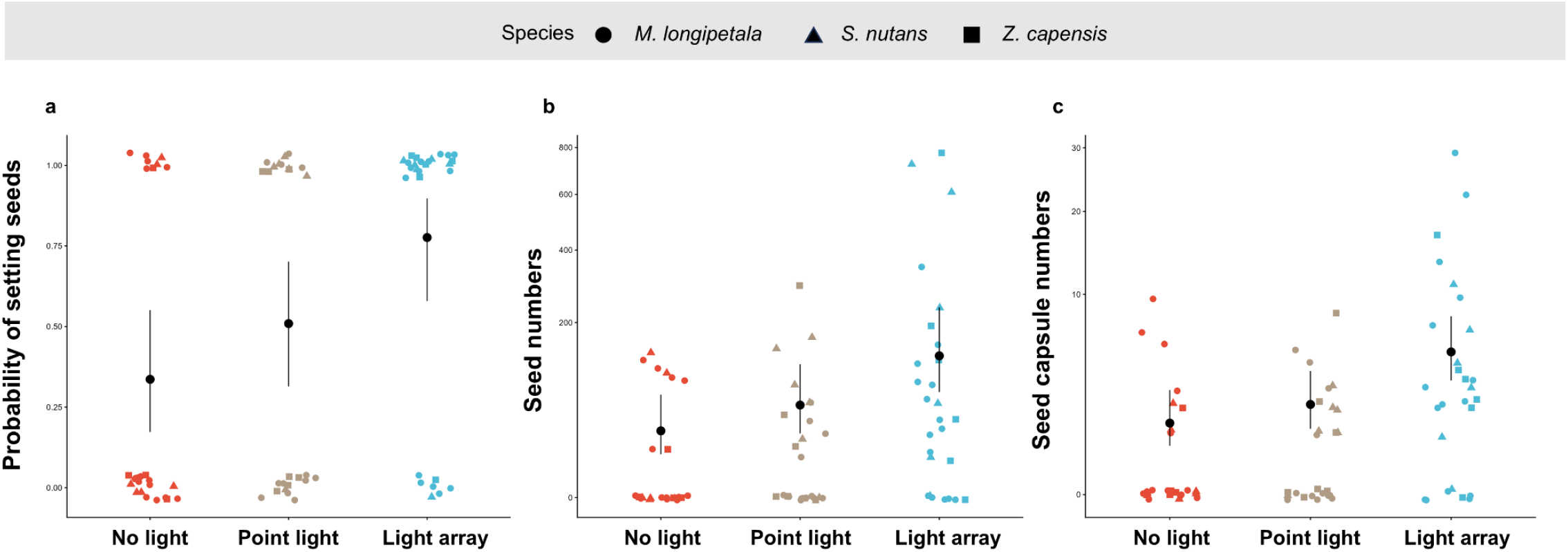
The effect of light configurations on the seed set of plant assemblages. **a**, Probability of seed set, defined as production of at least one seed by an individual plant. **b**, Seed number. **c**, Seed capsule number. Coloured points represent raw plant-level observations; shapes indicate species: *M. longipetala* (circles), *S. nutans* (triangles) and *Z. capensis* (squares). Black points and error bars show model-predicted means and 95% confidence intervals from retained models.

Experimental lighting also affected seed (χ^2^ = 14.9455, df = 2, *p* < 0.001, Fig. 2b) and seed capsule (χ^2^ = 14.5074, df = 2, *p* < 0.001, Fig. 2c) numbers. Relative to unlit controls, light arrays increased seed number (log-rate ratio = 1.51, 95% CI = [0.65, 2.72]) and seed capsule number (log-rate ratio = 1.38, 95% CI = [0.65, 2.39]). *S. nutans* produced more seeds (log-rate ratio = 0.88, 95% CI = [0.07, 1.85]) than *M. longipetala*, whereas *Z. capensis* did not differ in seed capsule production.

## 4. Discussion

The purpose of this study was to provide a small-scale field test of whether the spatial configuration of low-intensity artificial lighting can influence pollination success in moth-pollinated plants. Our results provide preliminary evidence that the spatial configuration of artificial lighting may be an overlooked factor shaping pollination responses to artificial light at night (ALAN). Low intensity light arrays increased the probability of seed set and enhanced seed and seed capsule production relative to unlit controls, whereas isolated point lights showed no clear effect. This suggests that ALAN may influence nocturnal pollination not only through the presence of illumination, but also through the spatial configuration of light sources.

Three limitations prevent us from drawing broader mechanistic conclusions. First, we did not directly observe or capture pollinators visiting the phytometer plant assemblages. We therefore cannot determine whether the observed reproductive responses were caused by changes in moth abundance, visitation rate or pollination efficiency. Second, although the phytometer plants used here are primarily associated with moth pollination, we cannot exclude contributions from other pollinator groups, by distinguishing between diurnal and nocturnal pollination (Macgregor et al., 2019). Finally, our lighting treatments were designed to represent contrasting field-realistic configurations rather than to equalize total light output. Consequently, the array treatment differed from isolated point lights in both spatial arrangement and illuminated extent. Equalizing total light output by clustering multiple bulbs at point-light locations would have produced larger and brighter illuminated patches, rather than true isolated point sources, and would therefore have represented a different ecological contrast. Future studies combining phytometer experiments with direct observation, camera monitoring or insect radar tracking (Degen et al., 2024), while experimentally separating light arrangement from total illumination, would help identify the behavioural mechanisms linking lighting configuration to plant reproductive success.

We found a positive effect of light arrays on the pollination success of moth-pollinated plants, contrasting with the common expectation that artificial light at night disrupts nocturnal pollinator activity and reduces pollination (Knop et al., 2017; Macgregor et al., 2017). One possible explanation is that arrays of lights altered moth movement across a larger area than isolated point lights, increasing encounter rates between moth pollinators and phytometer plants. However, because moth availability or movements were not directly observed, this mechanism remains hypothetical. Importantly, our experiment differed from many previous studies (Knop et al., 2017; Macgregor et al., 2017) in what we used low-intensity garden lights. The positive response to arrays may therefore reflect how multiple weak light sources collectively modified the local light environment. Positive effects of lighting on plant reproduction have also been reported by Macgregor et al. (2019), where the plant *Silene latifolia* had higher pollination success under full-night lighting than under unlit controls. These findings suggest that ALAN may not simply suppress pollination, but may reorganize nocturnal pollinator movement and redistribute pollination services across landscapes, allowing some plant species to benefit while others may be disadvantaged (Murphy et al., 2022).

In contrast to light arrays, point-light treatments did not increase seed set probability nor reproductive output relative to unlit controls. This suggests that isolated lights may not necessarily alter effective pollination, even if they potentially affect moth movement. This could be due to that point lights affect moth behaviour only within a restricted zone (Degen et al., 2016), or concentrate activity near the light source without increasing visits to nearby flowers. Alternatively, the low-intensity garden lights used in our experiment may have been insufficient to attract enough moth pollinators to alter plant reproductive output (Bennie et al., 2016; Briolat et al., 2021). Nevertheless, the absence of a point-light effect supports the idea that pollination responses to ALAN depend on lighting configuration, not simply the presence of illumination.

In summary, our results provide preliminary empirical evidence that the spatial arrangement of artificial lighting may influence the pollination success of moth-pollinated plants. Rather than acting simply through the presence or absence of illumination, the effects of ALAN may depend on how light sources are distributed across space. In fragmented and increasingly urbanized landscapes, low-intensity lighting (< 5 lumens) networks could potentially alter nocturnal pollinator movement among floral resources, producing corridor-like or redistribution effects under some conditions. Future experiments that equalize total light availability while varying spatial arrangement, combined with direct moth tracking, are needed to test whether artificial light arrays facilitate, redirect or restrict nocturnal pollinator dispersal.

## Acknowledgement

We thank Alanna Kelly for help with growing and maintaining the plants used in the experiment. This work was supported by the China Scholarship Council (Grant no. 20186190011).

## Data availability statement

Data are available to access in GitHub (https://github.com/Dongboli/experimental-data) for peer-review, and will make it available upon publish, using figshare repository.

