## Supplementary material for "Pollination responses to artificial light at night may vary with lighting arrangement": Supp. Info.

**List of content**:

**Figure S1 -** Emission spectra of the lights used in the experimental light arrays

**Figure S2 -** Experimental light arrays at night

**Figure S3 -** Species-specific sampling round of three phytometer species

**Table S1** - Kruskal–Wallis test on the starting density of flower bud

**Table S2** - The effect of lighting treatment and species on the probability and production of seed set.

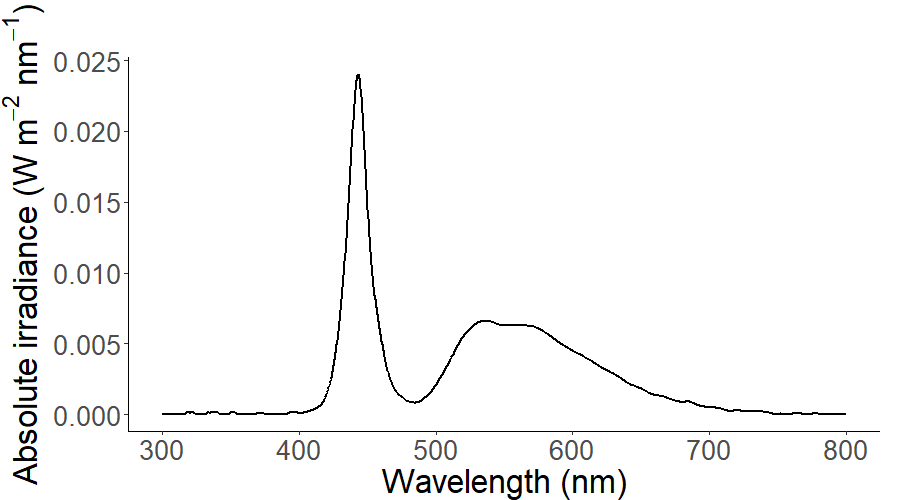

**Figure S1** – The emission spectra of the lights used in the experimental light arrays to test the effect of lighting configuration on pollination success in moth-pollinated plants. Irradiance displayed is an average of three measurements taken using a NIST calibrated StellarNet BLUE-wave spectrometer and CR2 cosine corrector (StellarNET Inc.).

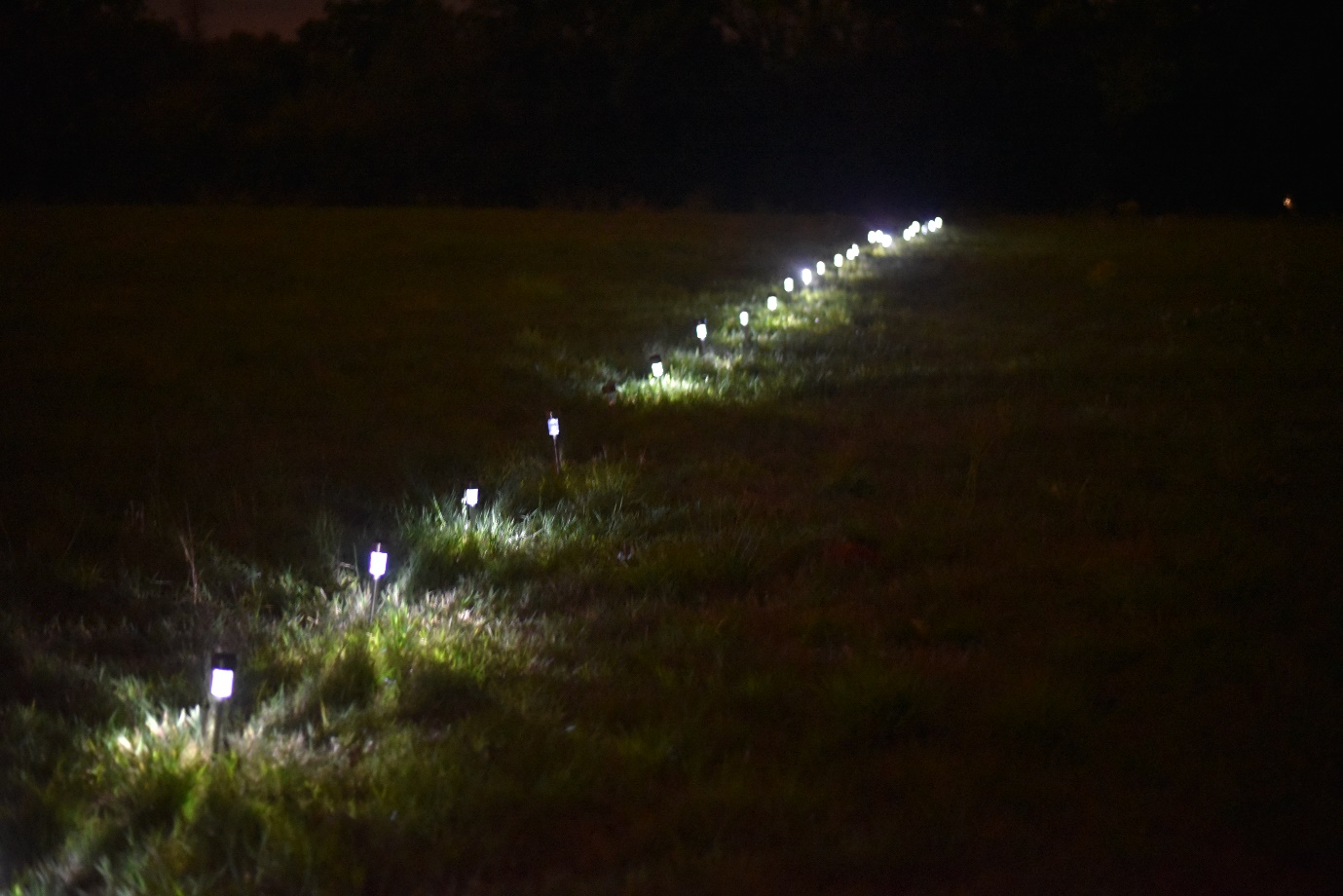

**Figure S2** – Experimental light arrays at night. Each replicate of light array treatment contained a linear array of 26 LED lights: one light at each endpoint and 24 additional lights spaced at 1 m intervals between them. Lights were solar-charged LED bollard garden lights (~5 lumen, 1.2 W), positioned approximately 20 cm above ground level and automatically illuminated after dusk for up to 8 h per night.

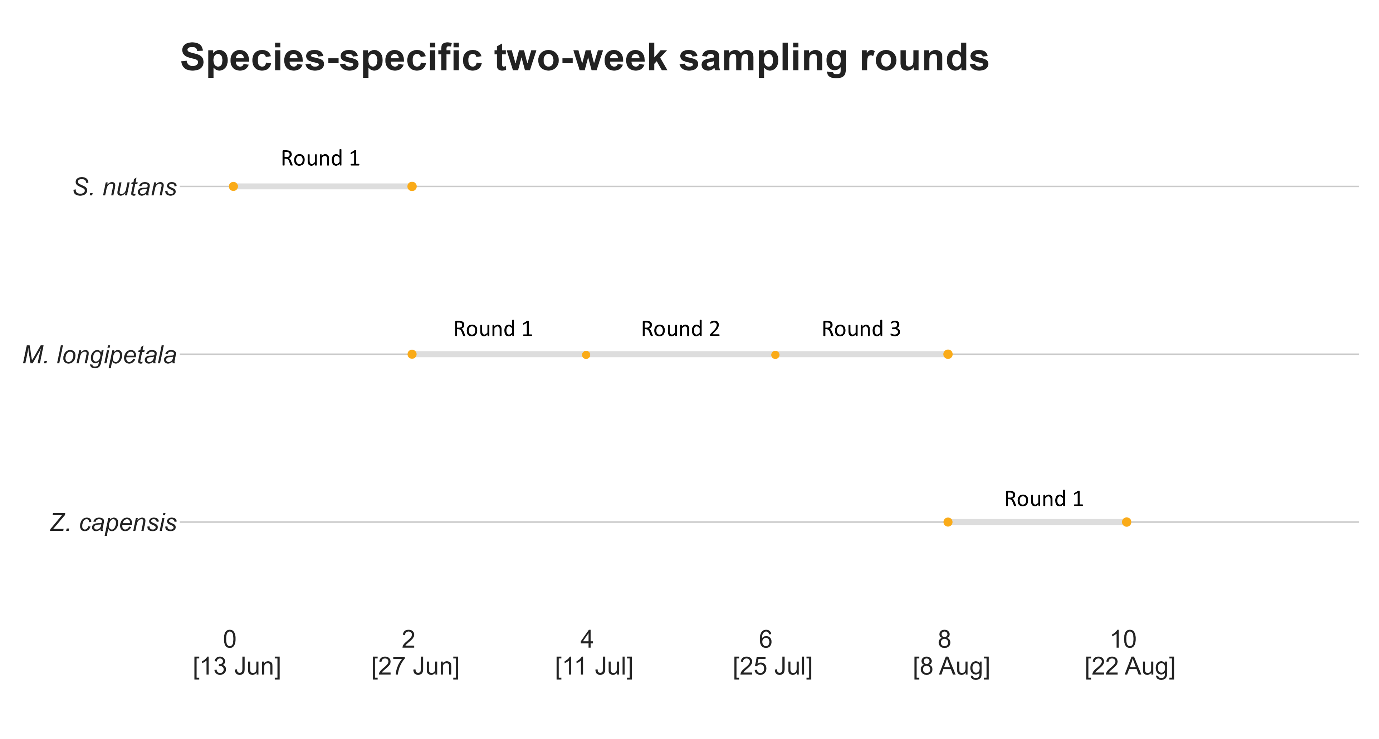

**Figure S3:** Species-specific sampling round of three phytometer species**:** *Matthiola longipetala* was deployed three two-week rounds, whereas *Silene nutans* and *Zaluzianskya capensis* were deployed one two-week rounds.

**Table S1** Non-parametric Kruskal–Wallis test on the effect of light treatment on the starting density of flower buds for each of the three phytometer species: *Matthiola longipetala*, *Silene nutans*, and *Zaluzianskya capensis.*

| Species | χ2 | df | *p* |
| --- | --- | --- | --- |
| *M. longipetala* | 1.8456 | 2 | 0.3974 |
| *S. nutans* | 2.5547 | 2 | 0.2788 |
| *Z. capensis* | 0.47045 | 2 | 0.7904 |

**Table S2** The effect of lighting treatment and species on the probability and production of seed set. Significant effects in the best fitted model (with lower AIC) were denoted bold.

| Response | Model structure | AIC | Variable | χ2 | df | *p* |
| --- | --- | --- | --- | --- | --- | --- |
| Probability | ~ Treatment + Species | **104.6867** | Treatment | 10.7639 | 2 | **0.005** |
|  |  |  | Species | 2.9304 | 2 | 0.231 |
|  | ~ Treatment * Species | 110.1643 | Treatment | 4.7814 | 2 | 0.092 |
|  |  |  | Species | 0.0850 | 2 | 0.958 |
|  |  |  | Treatment:  Species | 2.5224 | 4 | 0.641 |
| Seed number | ~ Treatment + Species | **593.7716** | Treatment | 14.9455 | 2 | **<0.001** |
|  |  |  | Species | 6.1611 | 2 | **0.046** |
|  | ~ Treatment * Species | 598.6626 | Treatment | 6.4083 | 2 | 0.041 |
|  |  |  | Species | 0.1657 | 2 | 0.921 |
|  |  |  | Treatment:  Species | 3.1734 | 4 | 0.529 |
| Seed capsule number | ~ Treatment + Species | **309.4947** | Treatment | 14.5074 | 2 | **<0.001** |
|  |  |  | Species | 0.8144 | 2 | 0.666 |
|  | ~ Treatment * Species | 316.0043 | Treatment | 7.9227 | 2 | 0.019 |
|  |  |  | Species | 0.1411 | 2 | 0.932 |
|  |  |  | Treatment:  Species | 1.4884 | 4 | 0.829 |
